# The Diversity and Evolution of Microbial Dissimilatory Phosphite Oxidation

**DOI:** 10.1101/2020.12.28.424620

**Authors:** Sophia D. Ewens, Alexa F. S. Gomberg, Tyler P. Barnum, Mikayla A. Borton, Hans K. Carlson, Kelly C. Wrighton, John D. Coates

## Abstract

Phosphite is the most energetically favorable chemotrophic electron donor known, with a half-cell potential (E°’) of −650 mV for the PO_4_ ^3-^/PO_3_ ^3-^ couple. Since the discovery of microbial dissimilatory phosphite oxidation (DPO) in 2000, the environmental distribution, evolution, and diversity of DPO microorganisms (DPOM) has remained enigmatic and only two species have been identified. Here metagenomic sequencing of phosphite enriched microbial communities enabled the reconstruction and metabolic characterization of 21 novel DPOM. These DPOM spanned six classes of bacteria, including the *Negativicutes, Desulfotomaculia, Synergistia, Syntrophia, Desulfobacteria* and *Desulfomonilia_A*. Comparing the DPO genes from the genomes of enriched organisms to over 17,000 publicly available metagenomes revealed the global existence of this metabolism in diverse anoxic environments, including wastewaters, sediments, and subsurface aquifers. Despite their newfound environmental and taxonomic diversity, metagenomic analyses suggested that the typical DPOM is a chemolithoautotroph that occupies low-oxygen environments and specializes in phosphite oxidation coupled to CO_2_ reduction. Phylogenetic analyses indicated that the DPO genes form a highly conserved cluster that likely has ancient origins predating the split of monoderm and diderm bacteria. By coupling microbial cultivation strategies with metagenomics, these studies highlighted the unsampled metabolic versatility latent in microbial communities. We have uncovered the unexpected prevalence, diversity, biochemical specialization, and ancient origins of a unique metabolism central to the redox cycling of phosphorus, a primary nutrient on earth.

**Significance Statement:** Geochemical models of the phosphorus (P) cycle uniquely ignore microbial redox transformations. Yet phosphite is a reduced P source that has been detected in several environments at concentrations that suggest a contemporary P redox cycle. Microbial dissimilatory phosphite oxidation (DPO) converts soluble phosphite into phosphate, and a false notion of rarity has limited our understanding of its diversity and environmental distribution. Here we demonstrate that DPO is an ancient energy metabolism hosted by taxonomically diverse, autotrophic bacteria that exist globally throughout anoxic environments. DPO microorganisms are therefore likely to have provided bioavailable phosphate and fixed carbon to anoxic ecosystems throughout Earth’s history and continue to do so in contemporary environments.

## Introduction

Phosphite (PO_3_ ^3-^) is a highly soluble, reduced compound that can account for over 30% of the total dissolved phosphorus in diverse environments^1,2^. Evidence suggests that meteorite impacts deposited substantial phosphite quantities on early Earth, but its abiotic oxidation to phosphate after the Great Oxidation Event (∼2.5 billion years ago [Gya]) is assumed to have rendered phosphite negligible in neoteric environments^3^. Surprisingly, phosphite has been detected in diverse reducing environments, and up to 1 µM was observed in some surface waters, suggesting contemporary neogenesis^1,3^. Geothermal and hydrothermal systems may generate phosphite through metal phosphide corrosion and iron-mediated phosphate reduction, and some phosphite may be derived from biological phosphonate degradation or anomalous phosphate reduction^1,4,5^. Meanwhile, some phosphite accumulation is likely attributable to anthropogenic activity because comparatively higher concentrations of phosphite have been identified in contaminated environments and industrial wastewaters^1,2,6^.

Despite its enigmatic distribution, functional gene presence in the IMG database^2^ predicts that phosphite is assimilated as a phosphorus source by approximately 1.5% of sequenced microorganisms^2,7–9^. However, the PO_4_ ^3-^/PO_3_ ^3-^ redox couple also has an extremely low potential (*E*^*o’*^= −650 mV), and microorganisms can alternatively use phosphite as a sole electron donor and energy source, excreting biogenic phosphate from cells^10^. With the low potential of the PO_4_ ^3-^ /PO_3_ ^3-^ redox couple, phosphite represents the most energetically favorable chemotrophic microbial electron donor described^11^, yet only two DPOM have been cultured, and only one has been isolated.

DPO was first identified in *Desulfotignum phosphitoxidans* strain FiPS-3, an autotrophic homoacetogenic facultative sulfate-reducing bacterium, isolated from Venetian brackish sediments^12^. DPO in FiPS-3 is attributed to the *ptx-ptd* gene cluster (*ptxDE-ptdCFGHI*), which FiPS-3 likely acquired through horizontal gene transfer (HGT)^13–15^. FiPS-3’s most closely related cultured isolate is incapable of DPO although the organisms share 99% 16S rRNA gene identity^16^. The second known DPOM, Ca. *Phosphitivorax anaerolimi* Phox-21, was enriched from wastewater collected in Oakland, California, and recently, another *Phosphitivorax* strain (Ca. *P. anaerolimi* F81) was identified in Danish wastewater^17,18^. Phox-21 grows chemolithoautotrophically with phosphite and carbon dioxide (CO_2_) as the sole electron donor and acceptor, respectively, and is the first naturally occurring species proposed to fix CO_2_ via the reductive glycine pathway^17,19,20^. The reductive glycine pathway has since been confirmed to naturally fix CO_2_ in wild-type *Desulfovibrio desulfuricans*^21^. Phox-21 harbors all *ptx-ptd* genes, but unlike FiPS-3, lacks *ptdG* (a putative transcriptional regulator) and shows no evidence of horizontal acquisition of the *ptx-ptd* cluster^2,13^. Understanding the evolutionary history of DPO metabolism is consequently limited by the existence of only two characterized DPOM whose *ptx-ptd* clusters exhibit deviating patterns of composition and inheritance.

Scarce representation also limits our understanding of the genes, organisms, and environments that support DPO. It is difficult to predict the range of DPO taxa because *D. phosphitoxidans* FiPS-3 and Ca. *P. anaerolimi* represent distinct taxonomic classes (*Desulfobacteria* and *Desulfomonilia_A*), and their closest relatives are either uncultured or unable to catalyze DPO^2^. The environmental context of DPO remains ambiguous since DPOM have only been identified in three distinct locations globally^16–18^. Furthermore, the *ptx-ptd* cluster has unresolved genetic diversity. *D. phosphitoxidans* FiPS-3 and Ca. *P. anaerolimi* species have *ptx-ptd* clusters with alternative synteny and gene composition, and the PtxD proteins from FiPS-3 and Phox-21 share only 55% amino acid sequence similarity^17^. Recognizing the breadth of hosts and environments supporting this metabolism and characterizing the underlying biochemistry and genetics would facilitate understanding of how DPOM impact the phosphorus cycle.

Here we present the selective enrichment of diverse DPOM in wastewater digester sludge from facilities around the San Francisco Bay area. Metagenome-assembled genomes (MAGs) uncovered 21 DPOM spanning three disparate phyla. Comparative genomics revealed conservation of energy generation and carbon utilization pathways among DPOM genomes, despite taxonomic diversity. We also identified DPO genes throughout global metagenome databases and described the diversity of the *ptx-ptd* cluster. The phylogeny of *ptx-ptd* genes suggests that DPO metabolism is vertically inherited as a conserved unit since before the split of monoderm (Gram-positive) and diderm (Gram-negative) bacteria. Collectively, our results show that DPO is widespread across diverse environments and bacterial taxa, and likely represents a vestige of ancient microbial life.

## Results

### Selective Enrichment

We hypothesized that DPOM are cultivatable from wastewater sludge because phosphite can represent up to 2.27% of total dissolved wastewater phosphorus^22^ and because both strains of Ca. *P. anaerolimi* were identified in wastewater digester sludge^17,18^. Accordingly, sludge from six San Francisco Bay area facilities were used to inoculate 30 enrichment cultures (Supplementary Dataset Table S1). All cultures were grown in bicarbonate-buffered basal medium amended with 10 mM phosphite and multivariate exogenous electron acceptors (CO_2_-only, CO_2_+SO_4_ ^2-^, or CO_2_+NO_3_ ^-^) (Supplementary Dataset Table S1). Rumen fluid (5% by volume) was added to stimulate DPOM growth^17^.

Phosphite oxidation was observed in 26 of 30 enrichments and across all six wastewater facilities (Fig. 1A & B, Supplementary Dataset Table S1). When stationary phase enrichments were re-spiked with phosphite, DPO activity resumed. No phosphite oxidation occurred in autoclaved controls (Fig. 1A). Based on prior experience^17^, the high percentage of active DPO enrichments was unpredicted, indicating a greater prevalence of DPOM than previously assumed.

**Figure 1:**
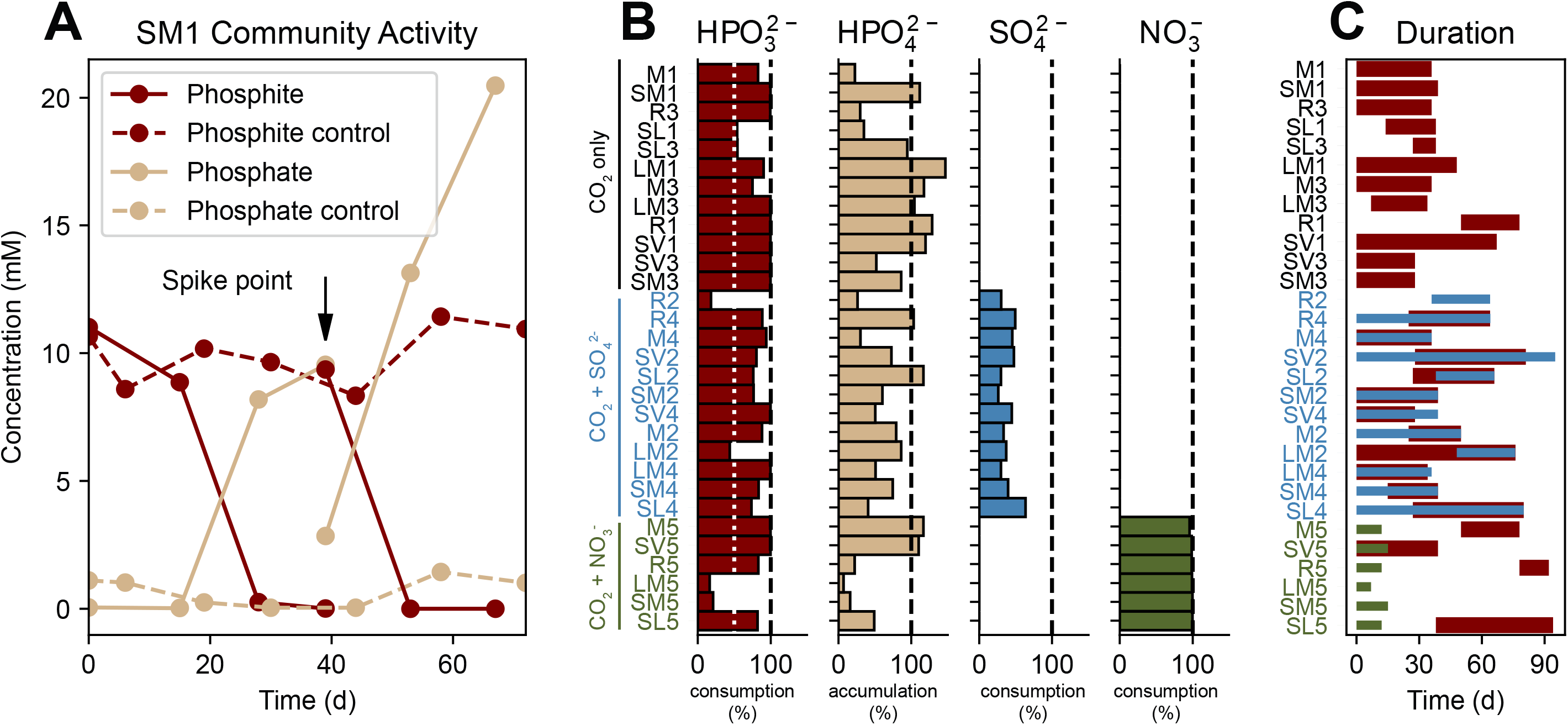
DPO Enrichment Activity. (A) Representative phosphite oxidation by the SM1 community. Temporal ion concentrations are shown for live (solid lines) or autoclaved (dashed lines) inoculum. Enrichments were amended with 10 mM phosphite at the spike point. (B) Percent change of measured ions for each enrichment community. Each row represents one community; each column displays the percent-accumulation or consumption of each titled ion. Row labels are colored according to the added electron acceptor (black, CO_2_ only; blue, CO_2_+SO_4_ ^2-^; green, CO_2_+NO_3_ ^-^). A white dotted line denotes 50% consumption of PO_3_ ^3-^. All percentages were calculated from concentration values *prior* to the first spike point. (C) Duration of ion depletion. Horizontal bars show the time frame for metabolic activity of each measured ion. Colors correspond with panel B (red, PO_3_ ^3-^; blue, SO_4_ ^2-^; green, NO_3_ ^-^).

### CO_2_ Preference

DPO was impacted by the amended electron-acceptor. Active enrichments with only CO_2_ supported the highest average phosphite oxidation rate (0.64±0.17 mM PO_3_ ^3-^/day for CO_2_ versus 0.56±0.10 and 0.50±0.20 mM PO_3_ ^3-^/day for NO_3_ ^-^ and SO_4_ ^2-^, respectively). CO_2_ also supported DPO from all six sample sites. Despite the availability of nitrate and sulfate, neither electron acceptor was definitively coupled to phosphite oxidation (Fig. 1C). While all amended cultures consumed nitrate, it was metabolized before phosphite oxidation was complete, suggesting utilization independent of DPO. In fact, when compared to other cultures with the same inoculum, nitrate delayed or even excluded DPO (Fig. 1C). Meanwhile, although sulfate was consistently consumed at the expected ratio, if reduced to sulfide coupled to phosphite oxidation (1 mol sulfate per 4 mols phosphite), the timing of sulfate consumption was variable and frequently offset from DPO (Fig. 1C). This suggests that sulfate reducers may be utilizing a reduced metabolite from DPO activity. Consistent with this, both of the characterized DPOM either grow preferentially (FiPS-3) or exclusively (Phox-21) by autotrophy and utilize CO_2_ as an electron acceptor. In the case of FiPS-3, the reduced carbon end-product is acetate^16^, which is readily utilized by sulfate reducers. Our results support a DPOM preference for CO_2_ and indicate that alternative electron acceptors may inhibit DPO activity^2^.

### DPOM Identification

To characterize the active DPOM, we recovered metagenome-assembled genomes (MAGs) from CO_2_-only enrichments. To identify candidate DPOM, we searched all MAGs using custom-built profile-HMMs (Files S1 – S7) for each of the seven *ptx-ptd* genes^13,15^. In total, 21 genomes had at least one gene from the *ptx-ptd* cluster (DPO MAGs), and of these, 19 were of high quality (>90% complete; <5% redundant) (Supplementary Dataset Table S3)^24^. DPO MAGs were enriched in all phosphite amended communities (compared to no-phosphite controls) (Fig. 2A) and were dominant in all but one community (SL1) (Fig. 2B). Furthermore, every sequenced community had at least one DPO MAG (Fig. 2B). These results confirmed that DPO activity in phosphite amended enrichments was dependent on the *ptx-ptd* genes and further indicates that these genes serve as effective probes for DPOM.

**Figure 2:**
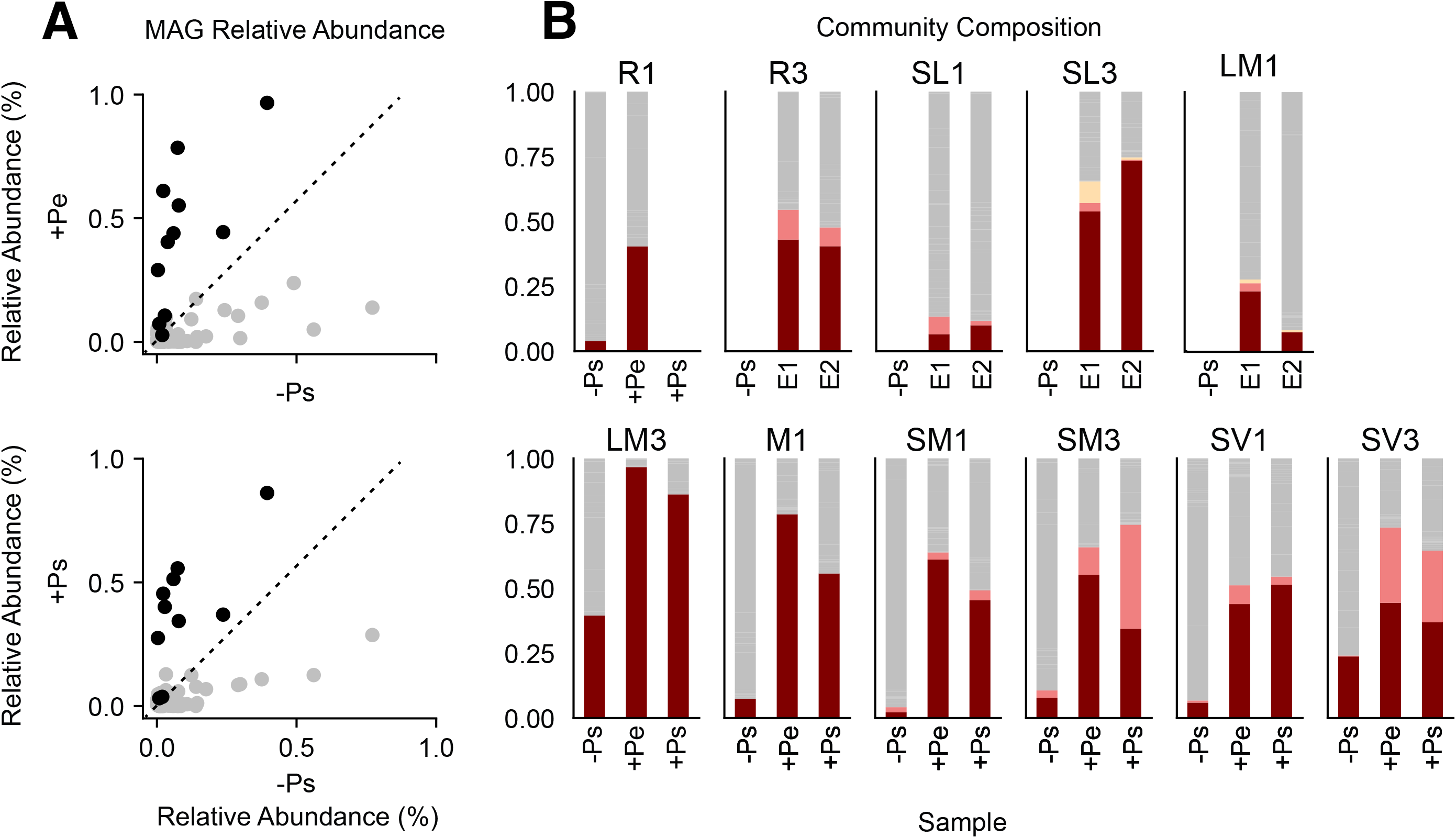
Relative Abundance of DPO MAGs. (A) Relative abundance of MAGs across samples. Each point represents one MAG. Color represents the presence (black) or absence (grey) of any *ptx-ptd* genes. Top panel compares samples from phosphite-amended exponential phase (+Pe) to no-phosphite (-Ps) controls. Bottom panel compares samples from phosphite-amended stationary phase (+Ps) to no-phosphite (-Ps) controls. (B) Relative abundance of MAGs across time. Each subplot represents one community, while each stacked bar represents the community composition of one sample. Colors indicate the dominant (maroon), second dominant (pink) and third dominant (yellow) DPO members, and all remaining community members (grey). Relative abundance was calculated by dividing the mean coverage of a single MAG by the sum of mean coverages for all MAGs in the respective sample.

### DPOM Taxonomy

DPOM taxonomy assignments were made using (i) reconstructed 16S rRNA gene fragments^25^, (ii) multigene alignments using the genome taxonomy database (GTDB)^26^, and (iii) alignment of the ribosomal S8 proteins (rpS8)^27^. Assignments were congruent in each instance and visualized in Figure 3. Prior to our study, DPOM had been identified as belonging to only two taxonomic classes of the *Desulfobacterota* phylum. In contrast, DPOM in our enrichments span the monoderm-diderm taxonomic boundaries and include three phyla (*Desulfobacterota, Firmicutes*, and *Synergistota*) and six classes (*Negativicutes, Desulfotomaculia, Synergistia, Syntrophia, Desulfobacteria* and *Desulfomonilia_A*) (Fig. 3).

**Figure 3:**
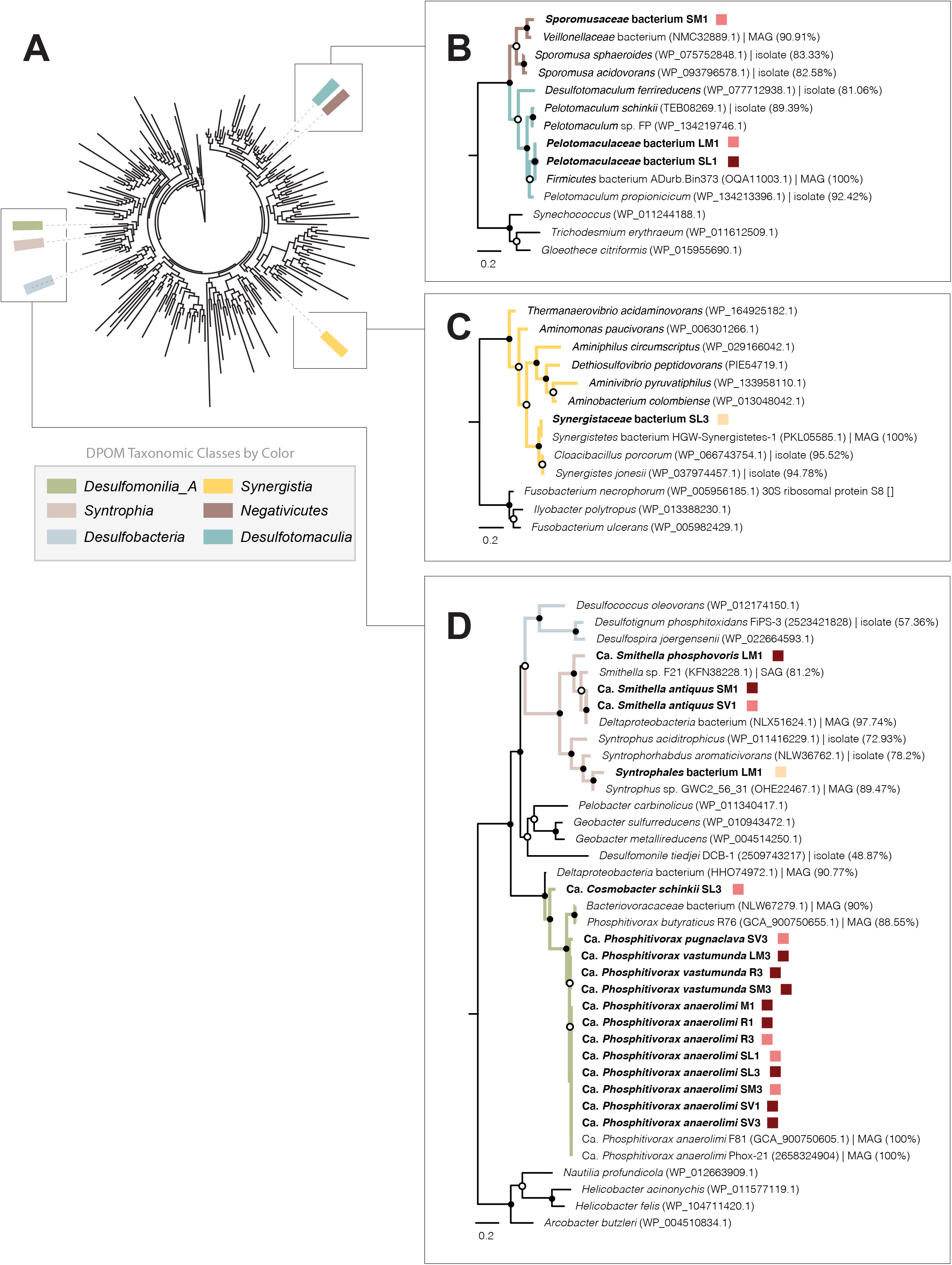
Phylogenetic Trees of DPO MAGs. A) A phylogenetic tree of bacterial genomes from the GTDB was visualized with AnnoTree^72^. Nodes of the tree represent class-level taxonomy, and those nodes with DPO organisms are highlighted according to the key. B-D) Phylogenetic trees of the rpS8 marker gene showing the relationship of DPO MAGs to their closest relatives. Panels depict DPO MAGs belonging to the same phyla: B) *Firmicutes*; C) *Synergistota*; D) *Desulfobacterota*. The DPO MAGs from this study are bolded. Colored squares represent their dominance rank from Fig. 2B. Each close relative is annotated with its species name, accession number and genome-source type (isolate vs. MAG), as well as its percent identity to the most closely related DPO MAG from this study. Clades are colored and labeled by taxonomic class. Internal nodes with bootstrap support of >90% are indicated by closed circles and those with support of >70% by open circles. Scale bars: 0.2 change per amino acid residue.

*Desulfomonilia_A* was the most sampled class of DPOM (Fig. 3), comprising 13 of 21 DPO MAGs. They were enriched from all six sample sites and were present in nine communities. Furthermore, they were the most relatively abundant DPO MAG (representing >85%) in each of eight communities, indicating a possible advantage under our enrichment conditions (Figs. 2B, 3). *Desulfomonilia_A* is an uncultured class that has recently been distinguished from the *Desulfomonilia* (https://gtdb.ecogenomic.org/). Consistent with this, the *Desulfomonilia* are represented by *Desulfomonile tiedjei*, which shares just 49% rpS8 sequence identity to the most closely related DPO MAG of *Desulfomonilia_A*^28,29^ (Fig. 3). All DPO MAGs of the *Desulfomonilia_A* class belong to the uncultured order UBA1062 (previously denoted GW-28)^17^, which includes Ca. *Phosphitivorax* (Supplementary Dataset Table S3). The monophyletic separation of the *Desulfomonilia_A* DPOM supports the hypothesis that Ca. *Phosphitivorax* species are part of a unique order, and possibly a unique class, for which DPO is a common metabolic feature^17^.

Beyond the *Desulfomonilia_A* DPOM, we recovered eight additional genomic representatives from four novel classes (*Negativicutes, Desulfotomaculia, Synergistia*, and *Syntrophia)* (Fig. 3). While most of these are minority DPOM in their respective communities, at least three (*Pelotomaculaceae* SL1, Ca. *Smithella* SM1, and Ca. *Smithella* LM1) dominate their DPOM populations (>83%) (Figs. 2B, 3). The Ca. *Negativicutes* and Ca. *Desulfotomaculia* MAGs represent the first DPO genomes taxonomically assigned to the *Firmicutes* phylum, highlighting the broad evolutionary divergence of DPOM^30,31^.

The closest cultured relatives of DPO MAGs share 57-95% rpS8 amino acid sequence identity (Fig. 3), which surpasses the species threshold (<98.3%)^27^. Furthermore, multigene classification by GTDB designates these related isolates as belonging to at least different genera^27^, making predictions about DPOM physiology from taxonomy unreliable. Regardless, all characterized DPO MAG relatives, regardless of taxonomy, are obligately anaerobic chemoorganotrophs. Furthermore, the *Desulfomonilia_A, Desulfotomaculia*, and *Syntrophia* classes contain canonical representatives that are dependent on syntrophic associations^18,31–34^. The phylogenetic relatedness of our DPOM to notoriously fastidious syntrophic organisms could explain the difficulty in isolating DPOM^17^.

The 16S rRNA gene is the canonical taxonomic marker for resolving microbial speciation. While not present in all DPO MAGs, 86% (n=18) contained the 16S rRNA gene, enabling refined taxonomic analyses (Supplementary Dataset Table S4). To capture the novelty of enriched DPOM, we used EMIRGE to reconstruct full-length 16S rRNA gene sequences that were BLAST searched in the SILVA database^25,35^. We determined that the DPOM represented 14 new strains, six new species, and one new genus based on standardized relatedness metrics^27^. Proposed names and etymologies are provided in Supplementary Dataset Table S4. The novel genus, represented by *Cosmobacter schinkii* SL3 (named in recognition of Bernhard Schink, for his exemplary contributions to microbiology and discovery of the first DPOM^12,16^), is the second characterized genus of the *Desulfomonilia_A* UBA1062 order, in addition to Ca. *Phosphitivorax*. Consequently, UBA1062 was expanded to include two genera and five species (Fig. 3).

### Metabolic Traits

The genomes of FiPS-3 and Phox-21 have been used to predict the mechanism for DPO energy conservation^13,17^. In the model (Fig. 4), the Ptx-Ptd protein cluster is hypothesized to oxidize phosphite and generate NADH and ATP through substrate level phosphorylation. Alternative reducing equivalents are likely produced via a Na^+^ motive force, ferredoxin, and an electron confurcation mechanism. The model proposes CO_2_ to be fixed into biomass via the reductive glycine pathway, as was suggested for Phox-21^17^. In contrast, FiPS-3 utilizes the Wood-Ljungdahl pathway^13^. By comparing the genomes of DPO MAGs to FiPS-3 and Phox-21, we found highly conserved metabolic traits beyond the *ptx-ptd* gene cluster, regardless of taxonomy.

**Figure 4:**
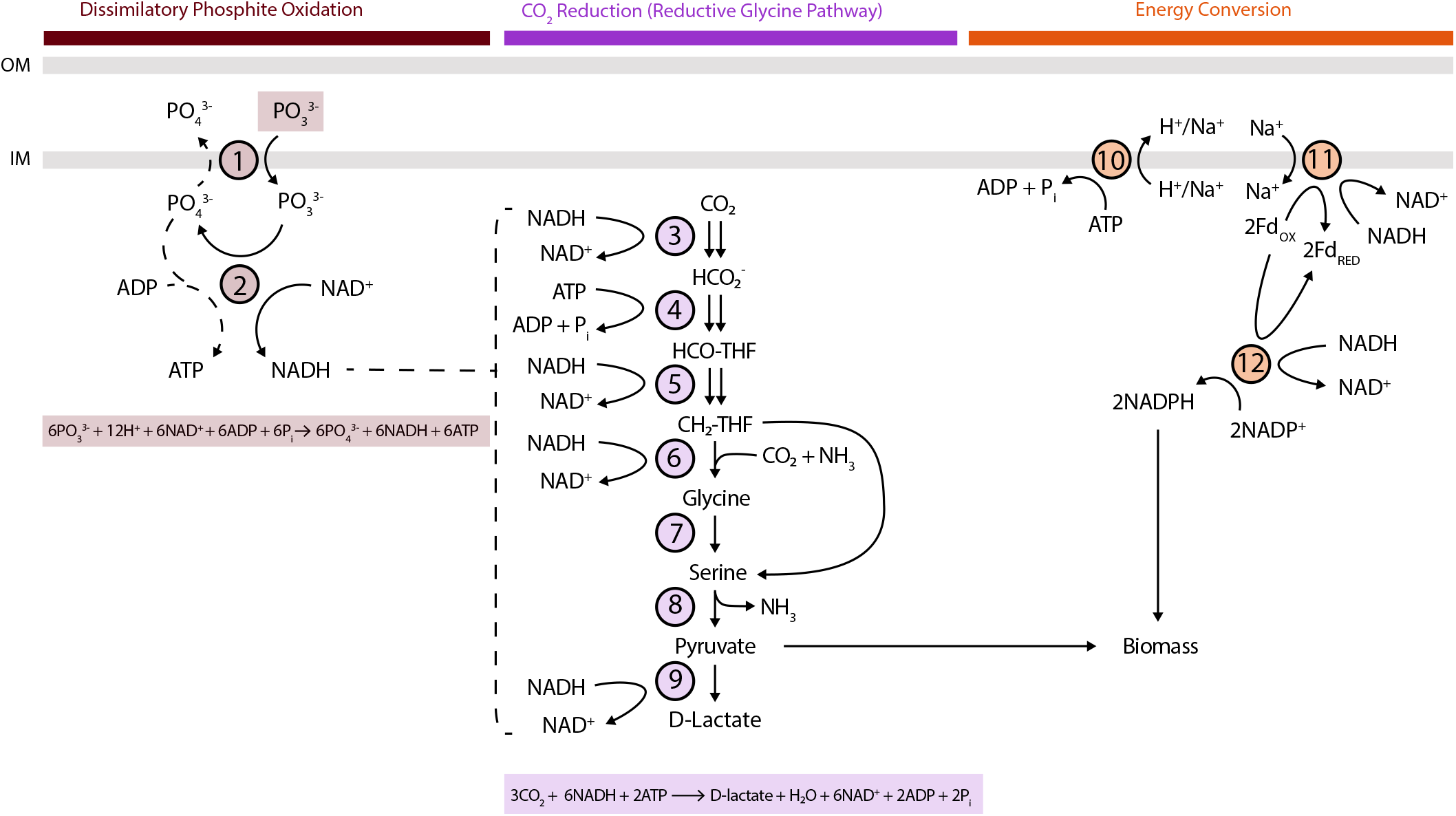
Metabolic model of energy conservation by *Desulfomonilia_A* DPOM. (adapted from Figueroa, *et al*.)^17^. Dotted lines represent mechanisms that have not been biochemically confirmed. Balanced equations are provided for phosphite oxidation and CO_2_ reduction to D-lactate. **Dissimilatory Phosphite Oxidation proteins**: (1) PtdC, phosphite-phosphate antiporter; (2) PtxDE-PtdFHI, putative phosphite dehydrogenase protein complex. **CO**_**2**_ **Reduction (Reductive Glycine Pathway) proteins**: (3) FdhAB/FdoGHI, formate dehydrogenase; (4) Fhs, formate:THF ligase; (5) FolD, methylene-THF dehydrogenase/methenyl-THF cyclohydrolase; (6) glycine cleavage system (GcvH, lipoyl-carrier protein; GcvPAB, glycine dehydrogenase; GcvT, aminomethyltransferase; Lpd, dihydrolipoyl dehydrogenase); (7) GlyA, serine hydroxymethyltransferase; (8) SdaA/IlvA, serine dehydratase/threonine dehydratase; (9) LdhA, D-lactate dehydrogenase. **Energy Conversion proteins:** (10) ATP synthase complex (11) Rnf, sodium-translocating ferredoxin:NAD oxidoreductase complex (12) NfnAB, NAD-dependent ferredoxin:NADP oxidoreductase.

### Energy Conservation

Like Phox-21, all DPO MAGs were missing a classical electron transport chain (ETC), as complexes II-IV were either absent or incomplete (Fig. 5). *Sporomusaceae* SM1 of the *Negativicutes* class had a complete NADH-quinone oxidoreductase (complex I), including the N, Q, and P-modules for NADH dehydrogenase activity, quinone reduction, and proton translocation, respectively. However, all other DPO MAGs only contained N-module subunits (Fig. 5, Supplementary Dataset Table S5). The N-module houses the FMN and FeS clusters for electron transport, as well as the NADH binding site. It also chimerically associates with other protein complexes, such as formate dehydrogenases, catalyzing reversible NADH-dependent formate production^36,37^. Poehlein *et al*. suggested that the FiPS-3 N-module may directly transfer electrons from NADH to ferredoxin^13^. However, direct NADH-dependent ferredoxin reduction is thermodynamically unfavorable^38^. Furthermore, the N-module of DPO MAGs is located in various genomic contexts, making it unclear whether the commonality is uniquely associated with DPO activity or with alternative cellular functions.

**Figure 5:**
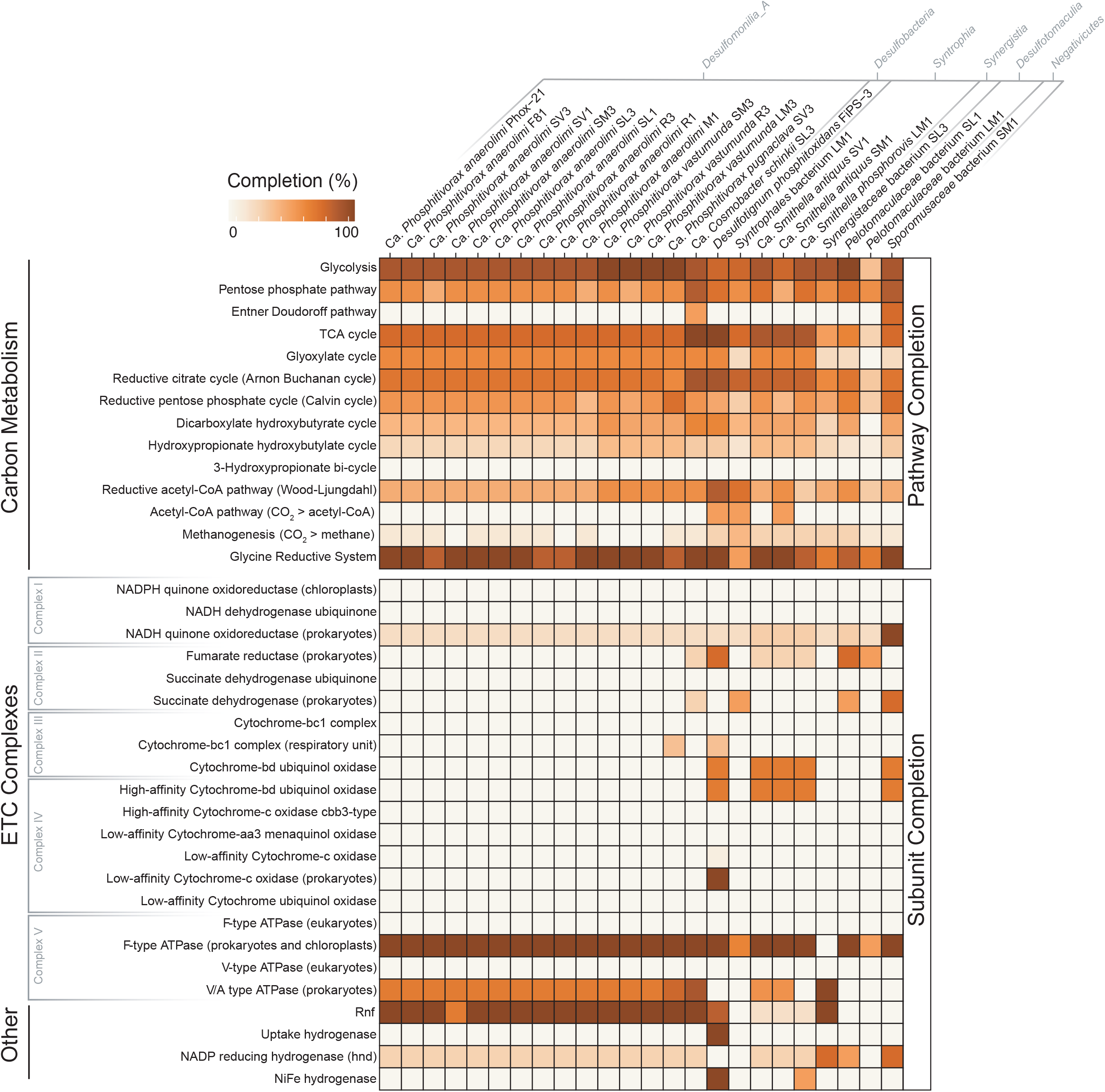
Carbon and Energy Metabolism of DPO MAGs. Each DPO MAG was subjected to metabolic analysis via DRAM^63,73^. Within this heatmap, each cell represents a metabolic pathway (columns) for each DPO genome (rows). The number of genes for a given pathway is described by percent completion ranging from 0% (white) to 100% (brown). Pathways are organized into modules related to carbon metabolism, electron transport chain (ETC) complexes, and other enzymes referenced in the text. Organisms are annotated with their taxonomic class.

In Phox-21, ferredoxin reduction by NADH is attributed to a sodium translocating ferrodoxin:NADH oxidoreductase (Rnf) driven by a Na^+^ motive force^17^ (Fig. 4). Consistent with Phox-21, an Rnf complex was present in the *Synergistia* and nearly all *Desulfomonilia_A* DPO MAGs (Fig. 5). In contrast, the Rnf was absent from the *Negativicutes, Desulfotomaculia*, and *Syntrophia* DPO MAGs, suggesting that it is dispensable or replaceable for DPO activity. The ion motive force for Rnf activity in Phox-21 is likely provided by a cation-translocating F-type ATPase at the expense of ATP (Fig. 4). The F-type ATPase was present in every DPO MAG, except one (*Synergistaceae* SL3) which had the V-type (Fig. 5). While two genomes (*Syntrophales* LM1 and *Pelotomaculaceae* LM1) were missing several ATPase subunits, these were only 61% and 69% complete (Supplementary Dataset Table S3)^24^. Given the universal absence of an ETC in DPO MAGs, the ATPases are likely involved in ATP hydrolysis with the concomitant generation of a cation motive force.

### CO_2_ as an Electron Acceptor

No DPO MAGs harbored functional pathways for methanogenesis or common respiratory pathways (oxygen, nitrate, or sulfate), which is similar to Phox-21 and consistent with the absence of ETC complexes (SI Appendix Fig. S2). Furthermore, CO_2_ was the only exogenous electron acceptor available to DPOM in sequenced cultures. Consistent with Phox-21^17^, a physiological survey of one of our enrichments showed that CO_2_ was necessary and sufficient to support phosphite oxidation and growth (Fig. 6). As observed in Phox-21, comparative genomics of DPO MAGs revealed a notable absence of any canonical CO_2_-reduction pathways (Fig. 5). While FiPS-3 can use CO_2_ as an electron acceptor by reducing it to acetate via the Wood-Ljungdahl pathway^16^, carbon reduction in Phox-21 was attributed to the reductive glycine pathway^17^. This is comprised of the methyl branch of the Wood-Ljungdahl pathway, combined with the glycine cleavage system, serine hydroxymethyltransferase, and serine deaminase to produce pyruvate as an anabolic intermediate^17,21^ (Fig. 4). The Phox-21 final product of CO_2_ reduction remains enigmatic, as the genes for pyruvate conversion to acetate (phosphotransacetylase and acetate kinase) are missing from the genome. Lactate is a possibility, as the genomes of Phox-21 and all other *Desulfomonilia_A* DPO MAGs contain D-lactate dehydrogenase, which converts pyruvate to lactate at the expense of NADH (Fig. 4). This is an energetically favorable reaction that accounts for all reducing equivalents produced via phosphite oxidation according to figure 4 and:

**Figure 6:**
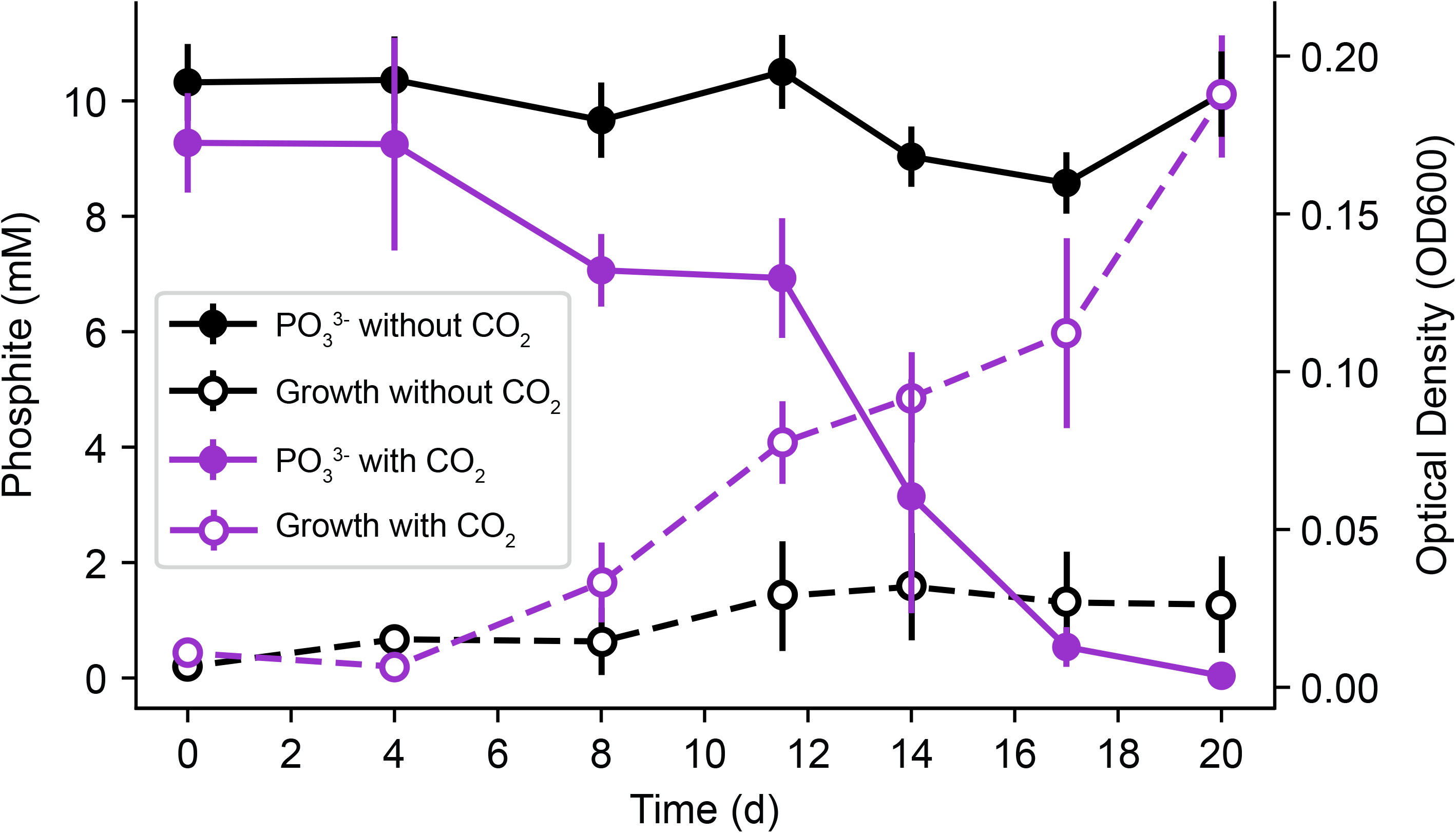
CO_2_ Dependent DPO Activity. Growth and phosphite concentrations were temporally monitored in the presence and absence of CO_2_ for the SV3 community. Autoclaved controls showed no activity. Error bars represent standard deviation of triplicate cultures.

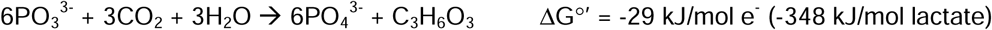

### CO_2_ Fixation to Biomass

In addition to serving as the electron acceptor for DPOM, CO_2_ is also fixed into biomass as the carbon source^17^ (Fig. 6). While none of the DPO MAGs contained any canonical CO_2_ fixation pathways^39^, twelve in the *Desulfomonilia_A, Negativicutes*, and *Syntrophia* classes had all the genes necessary for CO_2_-fixation to pyruvate via the reductive glycine pathway^21^ (Figs. 4 & 5). Of the residual nine DPO MAGs whose reductive glycine pathway was incomplete, four were missing homologs of serine deaminases, preventing the final conversion of serine to pyruvate (Supplementary Dataset Table S6). The remaining five DPO MAGs (ranging from 61.3 – 98.7% completion) were missing between one and four genes involved in formate and/or glycine transformations, severely impeding the overall pathway (Supplementary Dataset Tables S5, S6). It is possible that homologous enzymes may perform the reactions of missing genes, as might be the case for one genome (*Syntrophales* LM1) which harbored a serine-glyoxylate transaminase as opposed to the standard serine deaminase (Supplementary Dataset Table S6). Even if not a universal carbon fixation pathway in DPOM, our analyses suggest the reductive glycine pathway might be an important autotrophic mechanism across diverse DPO taxa. Carbon-tracing studies will be necessary to understand how individual DPOM use CO_2_ to simultaneously generate biomass and capture energy from phosphite oxidation.

### *ptx-ptd* Cluster Diversity

DPO activity in FiPS-3 and Phox-21 was attributed to the *ptx-ptd* gene cluster, and only organisms with *ptx-ptd* genes were enriched, positing this to be the dominant, or possibly sole, metabolic pathway underlying phosphite oxidation^13,14,17^ (Fig. 2A). To determine the prevalence and diversity of DPOM beyond our enrichments, we used the PtxD protein sequence from FiPS-3 as a marker gene to query the IMG/M protein sequence space (Supplementary Dataset Table S7). We recovered 15 positive hits that were phylogenetically compared to the PtxD from our enriched DPO MAGs and the two previously known DPO species (Fig. 7).

**Figure 7:**
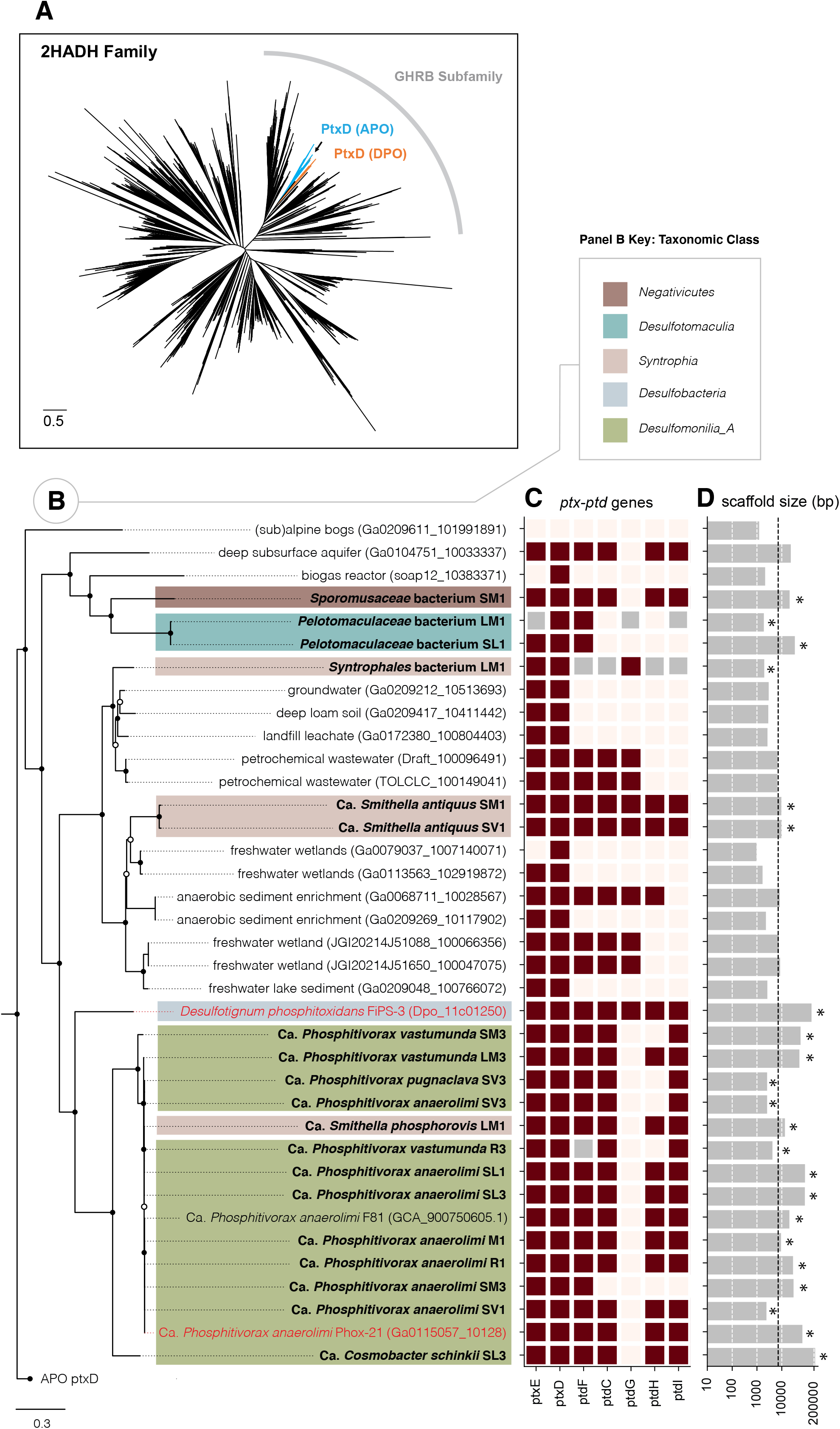
Phylogenetic tree of the phosphite dehydrogenase PtxD. A) The PtxD from IMG/M metagenomes and DPO MAGs were aligned with proteins from the 2-hydroxyacid dehydrogenase family (Pfam PF00389, set representative proteomes to 15%). Protein subfamilies were assigned based on Matelska et al.^40^. An arrow indicates the location of PtxD proteins that are associated with DPO PtdC but clade with assimilatory phosphite oxidation PtxD (APO). Scale bar: 0.5 change per amino acid residue. B) Refined tree of all PtxD within the DPO-PtxD clade. PtxD from the IMG/M are in light black font and labeled with their source environment and scaffold ID. PtxD from our enriched DPO MAGs are bolded and labeled with their bacterial host name. PtxD that belong to a binned organism are highlighted based on their taxonomic class. Published organisms with validated DPO activity are in red font. Only genes adhering to the IMG/M data usage policy are shown. Internal nodes with bootstrap support of >70% are indicated by closed circles and those with support of >50% by open circles. Scale bar: 0.2 change per amino acid residue. C) The presence (maroon) or absence (light pink) of *ptx-ptd* genes in each genome was determined using custom pHMM models. Genes that were absent from a DPO MAG but present in the assembly are in grey, where phylogeny, tanglegrams, and synteny were collectively used to predict the most likely host. D) Horizontal grey bars display the size (bp) of the contig on which each PtxD was found and are in logarithmic scale to visualize the full range of contig lengths. The black dotted line indicates the minimum length for all seven *ptx-ptd* genes to be present, based on FiPS-3 sequences (7137 bp). Asterisks signify contigs that were binned. bp, base pair.

Our analysis revealed that the DPO-PtxD form a monophyletic clade that included all validated DPOM (i.e. FiPS-3, Phox-21, and our enriched DPOM). The DPO-PtxD belonged to the glyoxylate/hydroxypyruvate reductase B (GHRB) protein sub-family of the D-2-hydroxyacid dehydrogenases (2HADH). The closest relatives of DPO-PtxD are the sugar dehydrogenases and the PtxD homologs involved in phosphorus assimilation^40^ (Fig. 7A, Supplementary Dataset Table S8). The DPO PtxD can be distinguished from closely related proteins based on the presence of nearby *ptd* genes (Fig. 7). The closest non-DPO homolog (Ga0209611_10199181) of the DPO-PtxD lacks the remaining *ptx-ptd* genes in the inclusion-matrix (Fig. 7C), demonstrating the specificity of our custom pHMMs (File S1 – S7).

The DPO PtxD was found exclusively in anoxic environments (Fig. 7B). The predicted failure of DPOM to occupy oxic environments, despite the thermodynamic favorability of DPO coupled to oxygen respiration (Δ *G*^*o’*^ = −283 kJ.mol^-1^ PO_3_^3-^), suggests that metabolic proteins may be oxygen sensitive. Alternatively, DPO metabolism may be dependent on the biochemical pathways of anaerobes. While DPOM appear to be common members of diverse anoxic environments, further analyses will be required to describe their relative abundance in natural habitats.

### Evolutionary History

The DPO evolutionary history was ascertained using (i) genomic features (ii) comparative taxonomic clustering, and (iii) syntenic conservation. Within the DPO-PtxD clade, proteins clustered based on host taxonomy, and the PtxD was distinguishable at the genus level (Fig. 7B). The only deviation from this pattern was Ca. *Smithella phosphorovis* LM1 of the *Syntrophia* class, which had a PtxD lineage consistent with Ca. *Phosphitivorax* species of the *Desulfomonilia_A* class (Fig. 7B). The *ptx-ptd* cluster from Ca. *S. phosphorovis* LM1 occurred on a single contig (13,378 bp) that hosted an IS91 family transposase. This contig had a sequencing depth (64.7x) three-fold that of the bin’s average coverage (19.4x), and the GC content (57.4%) was 3.5% higher than the host genome mean GC content (53.9%). Together these findings suggest that, like FiPS-3 ^16^, Ca. *S. phosphorovis* LM1 likely acquired its *ptx-ptd* genes through horizontal gene transfer (HGT). Consistent with this conclusion, the LM1 community assembly did not include taxonomic marker genes for Ca. *Phosphitivorax* species, and the assembly graph supported the binning results, precluding a different bin-assignment for this contig.

In contrast to FiPS-3 and Ca. *S. phosphorovis* LM1, most PtxD clustered according to host taxonomy, indicating that most DPOM likely acquired their PtxD via vertical inheritance (Fig. 7B). Similar taxonomic clustering occurred for the PtdC and PtdF, further suggesting that the *ptx-ptd* genes are inherited as a metabolic unit (SI Appendix Fig. S3). Tanglegram analyses facilitate a coarse approximation of topological similarity between gene phylogenies, where crossing lines (“tangles”) indicate alternative evolutionary histories^41^. Comparisons of the PtxD, PtdC, and PtdF exhibited zero tangling, supporting a linked evolutionary history (SI Appendix Fig. S4). Although the phylogenetic trees of individual DPO genes showed alternative branching patterns, this was expected, as genes with functional differences are subject to unique selective pressures.

Synteny provides an alternative metric to gauge the unison of *ptx-ptd* gene evolution because: (i) linked genes tend to maintain organization throughout evolutionary history, and (ii) closely related taxa show high genomic stability^42,43^. We found that the individual *ptx* and *ptd* genes were always codirectional in the order *ptxED* and *ptdCF(G)HI*, respectively (SI Appendix Fig. S5). However, the directionality between the *ptx* and *ptd* gene clusters was variable and syntenic variation formed four distinct groups (Groups I-IV, SI Appendix Fig. S5) that correlated with host taxonomy. Groups I and IV do not contain *ptdG*, suggesting it is nonessential (SI Appendix Fig. S5). While other genes were frequently missing from the *ptx-ptd* cluster, synteny analysis suggested this is due to fragmented contigs (Fig. 6C, SI Appendix Fig. S5). For example, *Synergistaceae* SL3 was identified as a DPOM in our enrichments, but our pHMM search failed to identify it’s PtxD (Fig. 7). Synteny suggested that the PtxD was truncated downstream of PtxE, which was confirmed by BLAST alignment (SI Appendix Fig. S5).

Searching metagenome databases with additional DPO genes would likely reveal other DPO contigs that were split from the PtxD. This was the case when we mined the IMG/M database for PtdC and identified five additional contigs with divergent PtxD phylogeny (Fig. 6A, SI Appendix Fig. S4). While these divergent genes may indicate further DPO diversity, their contigs showed non-canonical *ptx-ptd* neighborhoods and are not yet represented by validated DPO cultures. For those *ptx-ptd* clusters that confidently represent DPOM, the predominance of vertical transfer was collectively supported by genomic features, taxonomy, and synteny.

## Discussion

We used cultivation-based investigations coupled to high-resolution metagenomics to clarify many of the confounding factors that have precluded understanding of DPO. Results from our studies have expanded the known diversity of DPOM ten-fold (from 2 to 21 genomes). Notably, phosphorus redox cycling coupled to CO_2_ reduction appears to be the primary metabolic niche occupied by DPOM. Although DPO coupled to any known inorganic electron acceptor (oxygen, manganese, perchlorate, nitrate, iron, sulfate etc.) is thermodynamically favorable, DPOM genomes encode sparse electron transport machinery and are largely devoid of the enzymes required to reduce these ions. CO_2_ was the only exogenous electron acceptor provided to our sequenced enrichments, and physiological experiments demonstrated a CO_2_-dependency. Yet DPOM also lacked canonical carbon reduction or fixation pathways. The reductive glycine pathway was present in many DPOM and may support CO_2_ fixation, but the method by which CO_2_ is fixed by the remaining DPOM is unknown, as is the end product of CO_2_ reduction (e.g. ethanol or lactate), begging future metabolomic analyses.

The highly specialized metabolic repertoire of DPOM is analogous to that of syntrophs, corroborating the observation that DPOM frequently belong to known syntrophic taxa^44^. Thermodynamically, phosphite is too energetically favorable an electron donor to require a syntrophic partner, but such a co-dependency would explain their resistance to isolation^17^. *D. phosphitoxidans* FiPS-3 remains the only cultured isolate to date, yet we failed to cultivate any close relatives of FiPS-3 in our enrichments, despite otherwise representing much of the DPO diversity present in global metagenomes. Furthermore, we found that FiPS-3 is phenotypically and genotypically anomalous when compared to other DPOM. FiPS-3 exhibits greater metabolic versatility than typical DPOM, reducing sulfate, thiosulfate, and nitrate as electron acceptors in addition to CO_2_ ^13,16,17^. FiPS-3 is also one of only two examples by which the *ptx-ptd* genes were likely acquired via HGT, suggesting that DPO is not its primary energy metabolism. Future efforts to cultivate DPOM may consequently be informed by our DPO MAGs, whose metabolic features suggest a dependence on limited substrates and a potential requirement for microbial partnerships.

Our DPOM spanned six classes of three bacterial phyla (*Desulfobacterota, Firmicutes* and *Synergistota*). Such sparse representation across diverse taxa is typically indicative of broad-host-range HGT, but phylogenetic analyses of the *ptx-ptd* gene cluster showed that DPO metabolic gene evolution mirrored the host taxonomy. This indicates that vertical transfer is the predominant mechanism of inheritance. Small variations in synteny further support the correlation between gene order and taxonomy while also suggesting that *ptx-ptd* genes have coevolved as a metabolic unit specialized for DPO metabolism.

Given the diversity of DPOM lineages that likely inherited the *ptx-ptd* gene cluster vertically, it is tempting to speculate the biological timescale for when DPO metabolism originated. The last common node for all known DPOM suggests that DPOM arose before the divergence of monoderm and diderm bacteria^45^. Mapping the divergence of these clades to geological timescales suggests that DPOM ∼3.2 Gya ^46^, contemporaneously with anoxygenic photosynthesis and ∼0.8 Gya after the evolution of methanogenesis^46^. This is consistent with the suggestion that phosphite composed 40-67% of dissolved phosphorus species in Archaean oceans (>3.5 Gya)^47,48^. The half-life of oceanic phosphite under a reducing atmosphere is expected to be 0.1-10 billion years, which would have allowed phosphite persistence on early Earth, possibly supporting a robust chemolithoautotrophic DPO population.

One would expect such an ancient metabolism to be detected more broadly across all bacteria. However, oxygenation of Earth’s atmosphere since the great oxidation event (∼2.5 Gya) has likely depleted ancient natural phosphite reserves, as oxidizing radicals abiotically oxidize phosphite on geological timescales^3,49^. Phosphite would consequently be too rare for DPO in most contemporary environments, and lack of positive selection would promote widespread gene loss^50^. Yet pockets of phosphite (0.1 – 1.3 μM) exist in diverse contemporary environments, and phosphite oxidizing metabolisms still occur in various habitats on extant Earth^10,22,51,52^. Environmental metadata from global metagenomes identified DPOM in multiple anoxic environments that represent relics of ancient Earth (i.e. oil reservoirs, deep subsurface aquifers) and serve as potential examples of contemporary phosphite accumulation (i.e. wastewater sludge, freshwater wetlands). A number of environments evidently continue to support phosphorus redox cycling. By coupling DPO to primary production via an uncharacterized CO_2_ reduction pathway, DPOM likely play a unique ecological role in any environment they inhabit.

## Methods

### Growth Conditions and Sampling

Enrichment inocula were obtained from six wastewater treatment facilities in the San Francisco Bay area of California (Supplementary Dataset Table S1). Serum bottles (150ml volume) (Bellco, Vineland, NJ, USA) containing basal media (45mL) were each inoculated with sludge (5mL) and incubated at 37 °C. Anoxic medium was prepared by boiling under N_2_/CO_2_ (80:20, v/v) to remove dissolved O_2_, and dispensed under N_2_/CO_2_ (80:20, v/v) into anaerobic pressure tubes or serum bottles. These were capped with thick butyl rubber stoppers and sterilized by autoclaving (15 min at 121 °C). The basal medium was composed of (per 1 L of DI water): 5 g NaHCO_3_, 12 g HEPES buffer, 1 g NH_4_Cl, 0.5 g KCl, 1.5 g MgCl_2_, 0.15 g CaCl_2_ (2H_2_O), 0.5 g L-cysteine HCl and 10 mL each of vitamins and trace minerals^53^. Saline medium additionally contained 20 g/L NaCl. Salt solutions of Na_2_HPO_3,_ Na_2_SO_4,_ and NaNO_3_ (10mM) were added from sterile anoxic stocks as needed. Rumen fluid (Bar Diamond Inc, Parma, ID, USA), prepared by degassing (30 minutes with N_2_) and autoclaving (121 °C for 30 min), was added to the basal media as required. Heat killed controls were autoclaved at 121 °C for 1 h. Samples for DNA extraction were pelleted by 30 min centrifugation at 10,000 rcf and stored at −80 °C. Samples for ion determination were filtered and stored at 4 °C prior to ion chromatography (IC) using the method described previously^17^. Cell growth was measured as optical density at 600nm (OD_600_) using a Genesys™ 20 Visible Spectrophotometer (Thermo Scientific).

### Metagenomic Assembly, Binning, and Annotation

Sequenced communities were grown in triplicate cultures amended with 5% rumen fluid with or without 10 mM phosphite (SI Appendix Fig. S1). DNA was extracted from the no-phosphite triplicates in stationary phase (-Ps), and the 10 mM phosphite triplicates in exponential phase (+Pe) and stationary phase (+Ps) (SI Appendix Fig. S1). Community R1 failed to reach stationary phase and was only represented by samples -Ps and +Pe. Communities LM1, R3, SL1, and SL3 failed to reproduce activity and were instead sampled from two previously active enrichments (E1 and E2) (SI Appendix Fig. S1). DNA was extracted using the DNeasy PowerLyzer Microbial Kit (Qiagen) and sequenced with an Illumina HiSeq 4000 (150 bp paired-end reads) at the UC Berkeley Vincent J. Coates Genomics Sequencing Laboratory. Reads were trimmed and filtered using Sickle v1.33 (quality threshold value of 20)^54^. Gene-level taxonomy was assigned using Centrifuge v1.0.1-beta-27-g30e3f06ec3 ^55^. Reads for each of the 11 communities were combined and co-assembled using MEGAHIT v1.1.2 ^56^ using the meta-sensitive preset. Reads were mapped to assembled contigs using BWA-MEM v0.7.17 ^57^ with default parameters. Contigs over 1000 bp from each combined assembly were binned into individual genomes using Anvi’o v5.4.0 ^58^. Communities with < 30,000 contigs (LM3, M1, R1, SM1, SM3, SV1, SV3) were binned manually using patterns of hierarchical clustering, sequencing coverage, GC content, and gene-level taxonomic assignments. Communities with > 30,000 contigs (LM1, R3, SL1, SL3) were binned automatically using CONCOCT then manually refined with the Anvi’o graphical interface^59^. Quality of metagenome-assembled genomes (MAGs) was measured from lineage-specific, conserved, single-copy marker genes using the CheckM v1.0.18 lineage workflow^60^. The resulting 11 co-assemblies consisted of 1900 Mbp, 1.99 million contigs, and 574 draft genomes (Supplementary Dataset Table S2). Only draft genomes of medium quality or greater (>50% completion; <10% redundant)^24^ were subjected to further study, resulting in 239 metagenome-assembled genomes (MAGs) that represent 60% (647 Mbp) of the binned contigs (Supplementary Dataset Table S3). Open reading frames were predicted from selected genomes using Prodigal v2.6.3 ^61^ and assigned taxonomy using the Genome Taxonomy Database toolkit (GTDB-Tk)^26^, which placed MAGs into protein reference trees using concatenated SCG sets. Contigs of interest were functionally annotated with Prokka v1.14.6 ^62^.

The DPO MAGs were also annotated with DRAM^63^, a genome annotation tool that provides metabolic profiles for each input genome. For contigs of interest, these annotations were compared to Prokka v1.14.6 annotations^62^. More detailed DRAM analyses are provided in Shaffer & Borton *et al*.^63^. The raw annotations containing an inventory of all database annotations for every gene from each input genome are reported in Supplementary Table S6. From the raw annotations, DRAM then summarizes key metabolisms across the genomes, with SI Appendix Figure S2 showing the DRAM Product output. All code for DRAM is available on github: https://github.com/shafferm/DRAM.

### Identification of Metagenomic DPO Proteins

DPO proteins (PtxD, PtdC, PtdF) were identified from publicly available metagenomes. The largest metagenomes (representing 90% of proteins from each ecosystem category) in the JGI Integrated Microbial Genomes and Metagenomes (IMG/M) database were collected (n=17,888) on August 1, 2018 (Supplementary Dataset Table S7). Sequence data from the IMG/M database were produced by the US Department of Energy Joint Genome Institute (http://www.jgi.doe.gov/) in collaboration with the user community. The FiPS-3 PtxD, PtdC, and PtdF were searched against all proteins using BLASTP with bit score thresholds of 270, 300, and 250 respectively. Positive hits were aligned using MUSCLE v3.8.1551 ^64^ and constructed into an approximately maximum-likelihood phylogenetic tree using FastTree v2.1.11 ^65^ with 1000 bootstrap resamplings. DPO proteins were defined as (i) those that formed a phylogenetically distinct clade with proteins from experimentally validated DPOM (ii) were found on a contig near at least one other putative DPO gene and (iii) were at least 90% the length of their homolog protein in FiPS-3. Protein sequences from the identified *ptx-ptd* gene clusters were used to create profile Hidden Markov Models (pHMMs) for each of the PtxDE-PtdCFGHI proteins using HMMER v3.2.1 ^66,67^. These pHMMs are available as supplementary files (File S1-S7). Bit score thresholds for stringent *de novo* identification of DPO proteins were determined by a reciprocal pHMM search on a subset of the IMG/M database (Supplementary Dataset Table S9). To compare the evolutionary relationships between the PtxD, PtdC, and PtdF, members of the DPO clade were dereplicated with CD-HIT v4.8.1 ^68^ by clustering proteins with 100% sequence similarity and selecting the largest contig to represent each gene cluster in a simplified phylogenetic tree. Tanglegrams comparing PtxD to PtdC and PtdF were generated with Dendroscope v3.7.2 ^69^. Gene synteny was visualized with SimpleSynteny ^70^, where genes were identified with BLAST and annotated according to our custom pHMMs.

### Characterization of DPO Genomes

Annotated proteins from all MAGs were searched for known DPO genes (*ptxDE-ptdCFGHI*) with our custom pHMMs. MAGs were operationally considered capable of DPO if they included at least one gene from the *ptx-ptd* gene cluster. The *ptx-ptd* genes that were absent from MAGs were searched for in all remaining contigs of the respective community.

A pHMM for the rpS8 was obtained from Wu *et al*. and applied to all DPO MAGs^67,71^. The rpS8 gene has been shown to effectively represent whole-genome average nucleotide identity (ANI) values^27^ and was present once in each DPO MAG. Each rpS8 was BLAST searched against the NCBI GenBank database to identify the closest relative, closest isolated relative, and informative representatives for phylogenetic analysis. Identified close relatives corresponded to the multi-gene taxonomy assignments of the GTDB (Supplementary Dataset Table S4). Sequences were aligned using MUSCLE v3.8.1551 ^64^, and an approximately-maximum-likelihood phylogenetic tree was constructed with 1000 bootstrap resamplings using FastTree v2.1.11 ^69^. Trees were visualized using FigTree v1.4.4 (http://tree.bio.ed.ac.uk/software/figtree/).

The 16S rRNA gene for each community was reconstructed from metagenomic reads using default parameters in EMIRGE with 50 iterations^25^. Reconstructed genes were classified using SILVA^35^ and mapped back to the 16S rRNA gene fragments of DPO MAGs. The novelty of each DPO MAG was determined by the rank of closest relatives in the GTDB, NCBI (rpS8), and SILVA (16S rRNA gene) databases (Supplementary Dataset Table S4). A DPO MAG was considered novel at the specified rank (i.e. species, genus) based on the following thresholds: (i) GTDB, considered novel if there were no logged relatives for that rank; (ii) NCBI (rpS8), considered a novel species if the closest relative was <98.3% identity; (iii) SILVA (16S rRNA gene) considered a novel species if the closest relative shared <96.7% identity and a novel genus if the closest relative shared <94% identity^27^. The novelty of a DPO MAG was assigned based on the lowest resolved taxonomic rank between all searched databases.

### Data Deposition

All metagenomic reads, assemblies, and curated metagenome-assembled genomes (MAGs; quality metrics >50% complete and <10% redundant) are available through the NCBI BioProject accession ######.

## Acknowledgments

We thank A. Englebrekston, I. Figueroa, M. Silverberg, Y. Liu, C. Thrash, J. Taylor, and S. McDevitt for laboratory support and guidance on evolutionary analysis and sequencing. Wastewater sludge samples were generously provided by Judy Walker (San Leandro Water Treatment), Aloke Vaid (Veolia Water North America, Richmond), Bob Wandro & Robert Huffstutler (Silicon Valley Clean Water), Jan Guy & Pete Dallabetta (San Mateo Waste Water Treatment Plant), Nimisha Patel (Sewerage Agency of Southern Marin), and Jimmie Truesdell (City of Livermore Water Resources Department). Funding for phosphorus redox cycling is provided by the Energy & Biosciences Institute (Berkeley, CA) and the US Department of Energy Genomic Science Program to J.D.C. Independent funding to S.D.E through the EBI-Shell Fellowship was supported by Shell International Exploration and Production Inc.

