## Supplementary figures and images for "The Diversity and Evolution of Microbial Dissimilatory Phosphite Oxidation"

### Fig S1

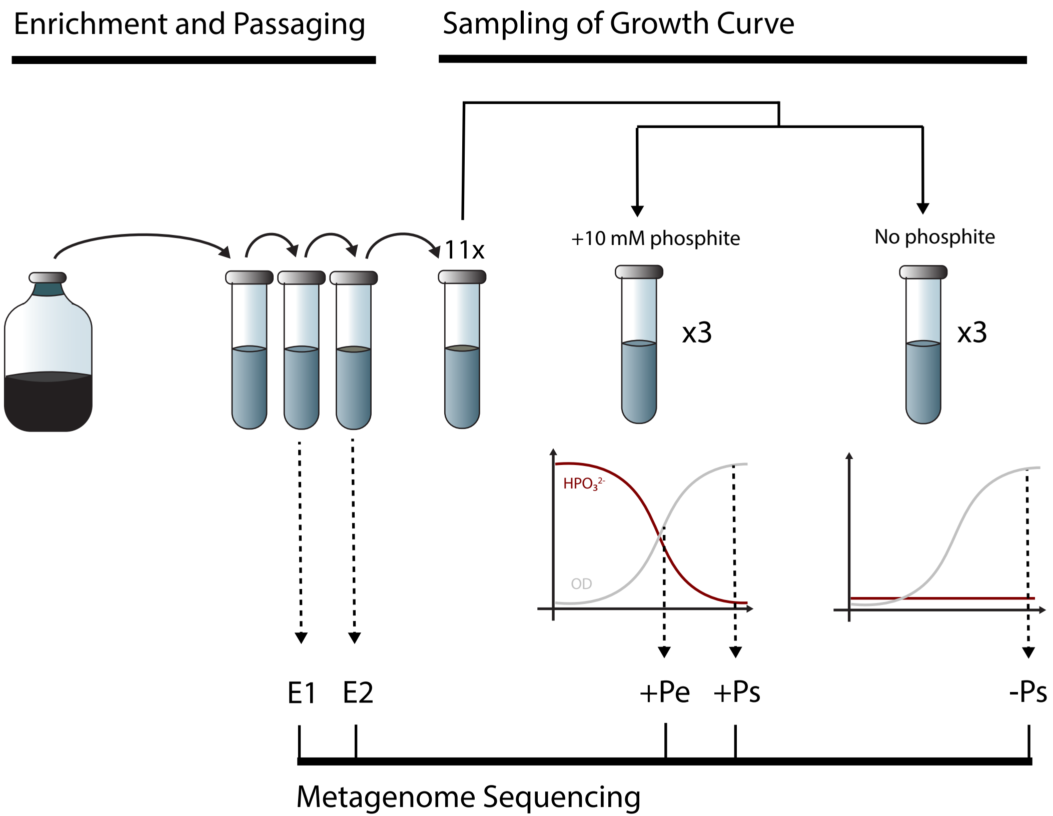

### Fig S2

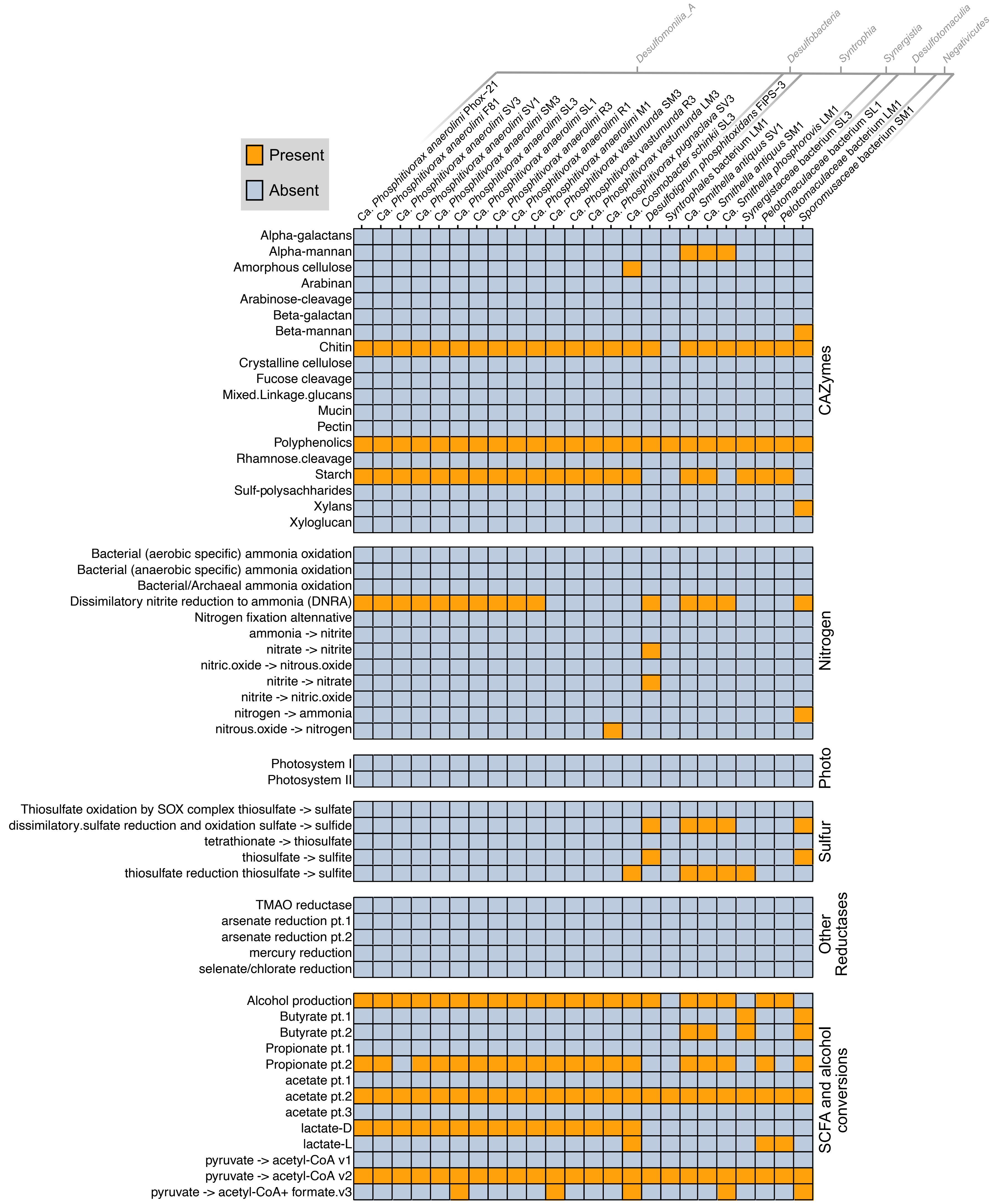

### Fig S3

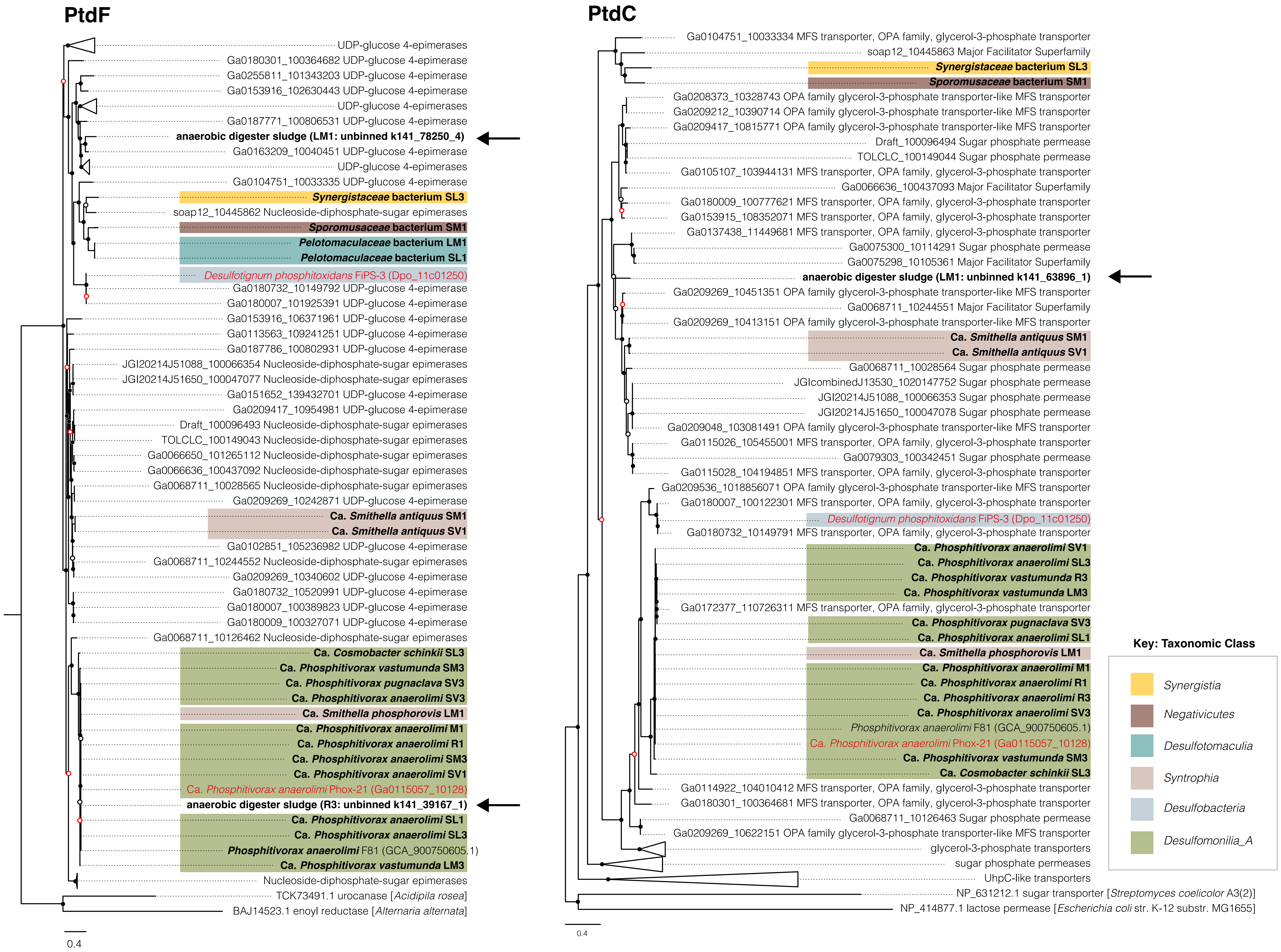

### Fig S4

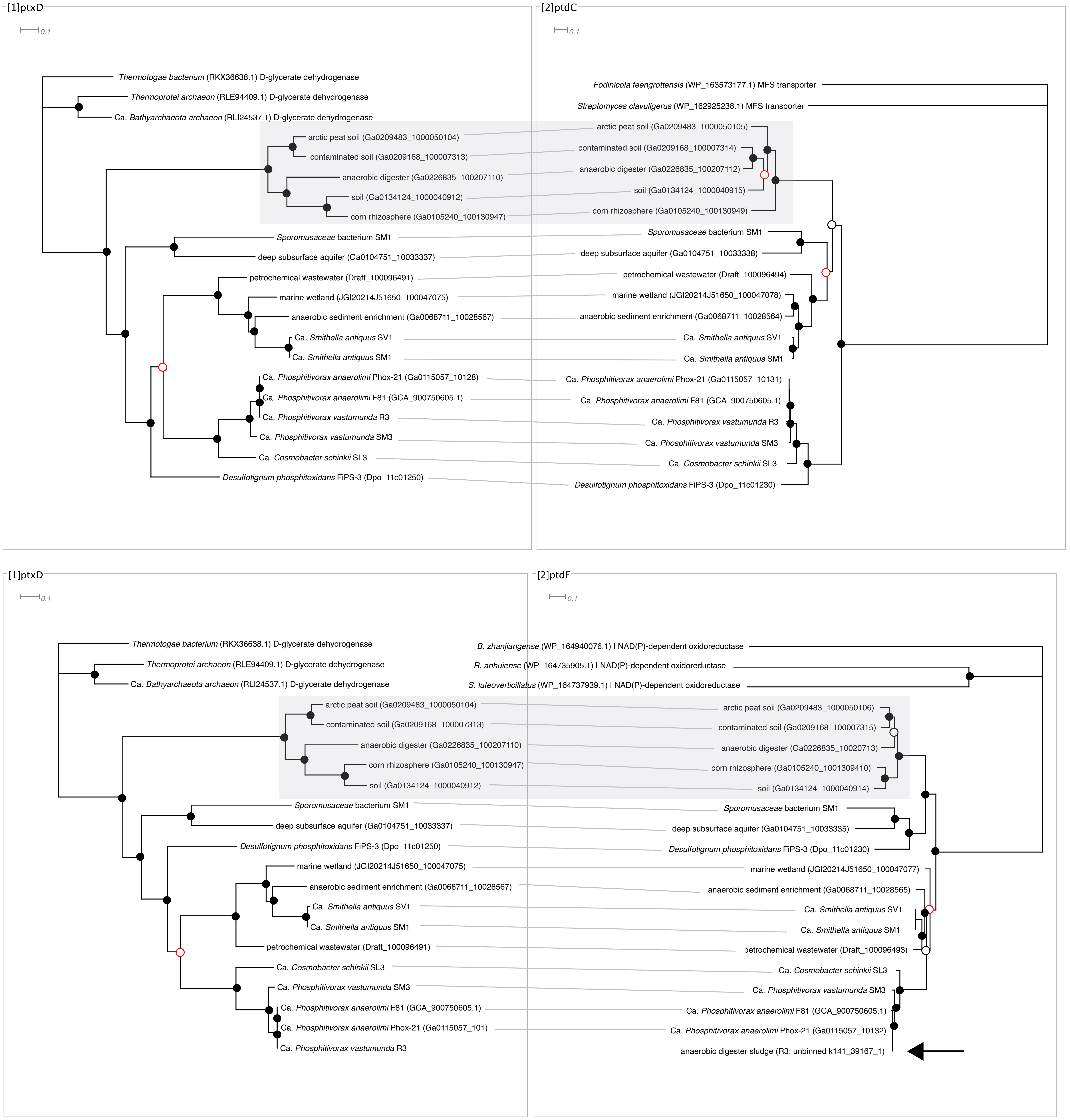

### Fig S5

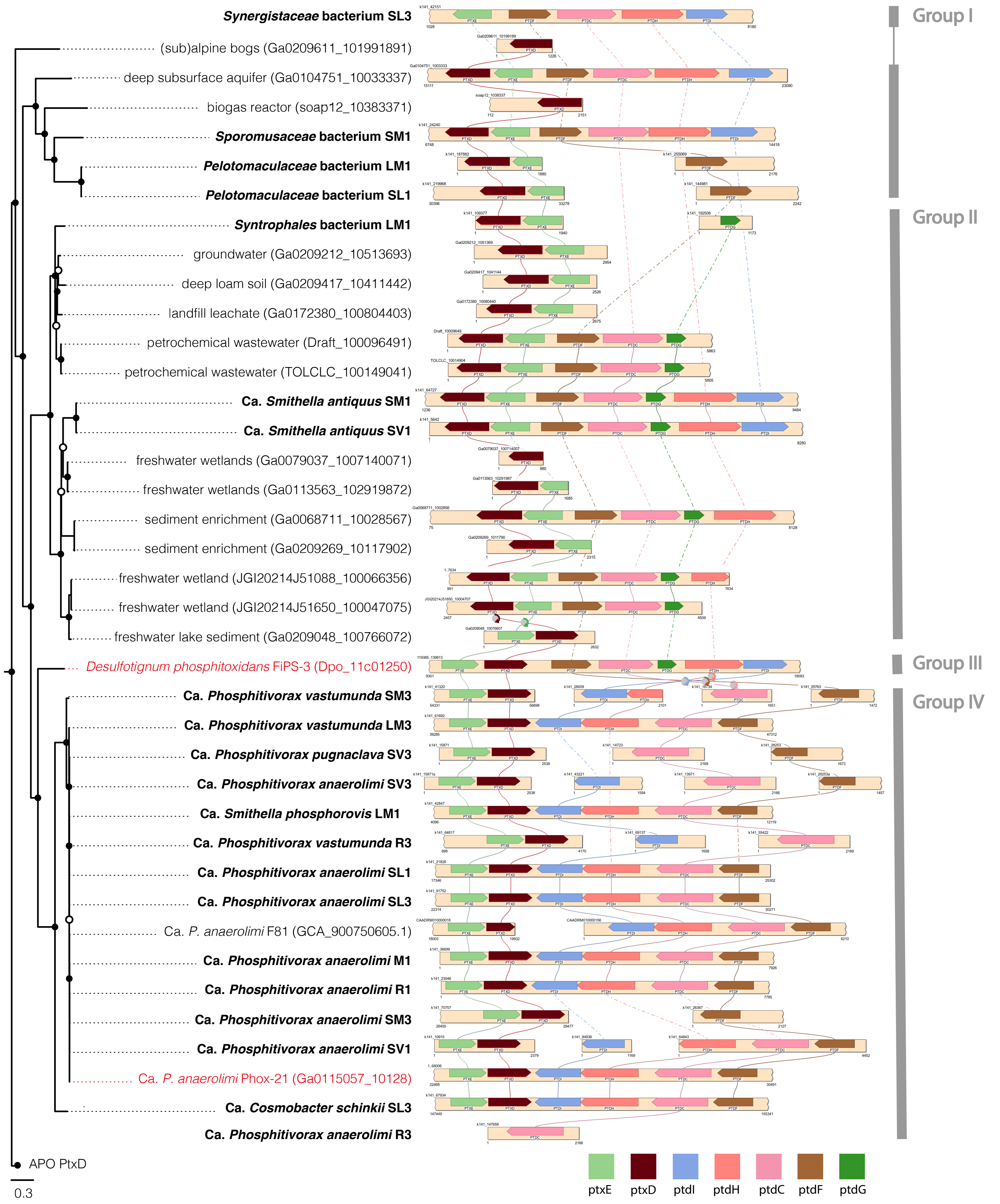
